# The genetic and social contributions to sex differences in lifespan in *Drosophila serrata*

**DOI:** 10.1101/2021.09.08.459506

**Authors:** Vikram P. Narayan, Alastair J. Wilson, Stephen F. Chenoweth

## Abstract

Sex differences in lifespan remain an intriguing puzzle for evolutionary biologists. A possible explanation for lower lifespan in males is the unconditional expression of recessive deleterious alleles in heterogametic X chromosomes in males (the unguarded X hypothesis). Empirical evidence, however, has yielded controversial results that can be attributed to differences in both genetic and social background. Here, we test the unguarded X hypothesis in *Drosophila serrata* using a factorial design to quantify the effects of genotype, sex, social environment, and their interactions on phenotypic variation for lifespan. Using an experimental approach, we manipulated two inbred laboratory genotypes and their reciprocal F1s, while controlling for different levels of density and mating status to account for any potential social effects. Our results also show subtle but significant genotype dependent effects for both density and mating, but ultimately find the unguarded X hypothesis insufficient to fully explain sexual dimorphism in *D. serrata* lifespan.

## Introduction

The question of why males and females differ in lifespan has long fascinated evolutionary biologists. While exceptions exist, across many taxa it is most often females that live longer than males (Austad, 2019). Despite a long history of ageing research, no proven or unifying theories have emerged, and studies still yield contradictory results. Sexual dimorphism in lifespan can arise in response to sex differences in selection on life histories. Males and females maximise reproductive fitness in different ways (Friberg, 2005, Maklakov et al., 2009) with males typically investing more in early reproduction than females, even at the cost of their own somatic maintenance and lifespan (Maklakov and Lummaa, 2013). Selection therefore alters the overall costs of reproduction for each sex, and affects the evolution of ageing by shaping sex-specific mortality rates (Promislow 2003; Bonduriansky et al. 2008). Sexual dimorphism in lifespan may also be caused by asymmetric inheritance of uneven numbers of sex chromosomes between males and females. This hypothesis posits that for species where males are the hemizygous sex, harmful recessive mutations on the X chromosome will always be expressed in males whereas they will commonly be masked by dominance in females (Trivers, 1985). A general prediction of this hypothesis coined the “unguarded X hypothesis” is that males should therefore on average have shorter lifespans than females.

Several studies have shown that variation in environmental or genetic background, can influence sexual dimorphism in lifespan (Kimber and Chippindale, 2013, Brengdahl et al., 2018b, Sultanova et al., 2018). Species of the genus Drosophila have featured prominently in aging research. In addition to *D. melanogaster* [see reviews by (Rogina, 2011) and (Piper and Partridge, 2018)], other species such as *D. simulans* (Ballard, 2005) have also been used as models for aging research. With the development of the *Drosophila serrata* Genome Reference Panel, a panel of re-sequenced lines (DsGRP) (Reddiex et al., 2018), *D. serrata* has now also emerged as a potential model for aging research. Here, we describe the results of a systematic analysis of lifespan comparisons in two highly inbred laboratory wild-type strains: DsGRP20 and DsGRP57. Using inbred lines can provide insight into how the underlying genetic architecture of lifespan varies in response to genetic and social conditions. For instance, Swindell and Bouzat (2006) showed that stressful environments such as increased competition and temperature had pronounced effects on mitigating lifespan reducing effects of inbreeding depression in *D. melanogaster*. While the existence of inbreeding depression on lifespan are well documented, how heterozygous and homozygous genotypes respond to social (Carazo et al., 2016, Sultanova et al., 2018, Brengdahl et al., 2018a) and environmental conditions (Tan et al., 2013, Brengdahl et al., 2018b, Sultanova and Carazo, 2019) such as mating and density is less well understood.

For *D. melanogaster*, the few studies where organismal condition was manipulated and lifespan was measured, both male- and female-biased effects on lifespan were found. This was true for genetic and environmental manipulations of condition. These studies highlight the importance of not just different genotypes, but also how sex differences in mating costs and behaviour affect survival rates (Burger and Promislow, 2004). This substantial variation in male and female responses emphasizes the importance of including not only both sexes, but also their social environment when analysing lifespan. Amongst the different social effects that have an impact on adult lifespan, mating activity and adult population density have been shown to influence longevity (Malick and Kidwell, 1966, Iliadi et al., 2009). In species of Drosophila, such as *D. virilis*, mating status significantly affected fly lifespan, with male and female virgins being affected very differently (Aigaki and Ohba, 1984). In *D. virillis* male sexual activity played the most important role amongst the complex interactions between both sexes. Mating status also affected the lifespan of both female and male *D. melanogaster* flies, though males were less affected (Koliada et al., 2020). The few systematic studies conducted on effects of high adult density, have found increased male sensitivity to variations in density, erratic mortality rates, and decreased mortality among higher density cohorts of middle-aged *D. melanogaster* females (Khazaeli et al., 1996).

This study aims to clarify how genetic background, sex, inbreeding, mating, and density act and interact with each other to shape lifespan in *Drosophila serrata*. In doing this we test the specific predictions of unguarded X and evaluate their sensitivity to genetic and social backgrounds. To quantify effects of genotype on lifespan we crossed fully inbred flies to generate outbred and reciprocal F1 flies (Vaiserman et al., 2013). To explore interactions with social contexts of mating (Aigaki and Ohba, 1984, Service, 1989, Zajitschek et al., 2013), we measured the lifespan of these flies as both virgins and non-virgins. Furthermore, we varied the population density of flies held together in a vial, as this is also known to affect lifespan and mortality rates (Graves and Mueller, 1993, Khazaeli et al., 1995, Khazaeli et al., 1996, Joshi and Mueller, 1997). This will ultimately bring us closer in our attempts to characterise sexual dimorphism in lifespan resulting from sex differences in selection as opposed to variation resulting from uneven numbers of sex chromosomes between males and females. We present evidence of genetic interactions with sex and also with mating and density on survival characteristics in *D. serrata*.

## Materials and Methods

### Fly stocks and culturing conditions

All analyses were carried out using fruit fly genotypes, DsGRP20 and DsGRP57, randomly chosen from the DsGRP (Reddiex et al., 2018). Flies were maintained in vials containing agar-sugar-yeast medium, in a temperature-controlled room at 25°C and a 12/12 h light/dark cycle. We then performed density-controlled crosses between the two lines to produce inbred (DsGRP20♂ × DsGRP20♀ and DsGRP57♂ × DsGRP57♀), and outbred reciprocal crosses (DsGRP20♂ × DsGRP57♀, and DsGRP57♂ × DsGRP20♀) from here on referred to as genotypes. All experimental flies were collected as virgins within 6h after eclosion, and male and female offspring from each cross were randomly allocated into the experimental treatments in a factorial design including the effects of mating, and density. Flies in the mated treatment were allowed to mate for 2 days, collected using CO2, sorted by sex, and transferred to experimental vials for the lifespan trial. For each cross, virgin and mated treatments were maintained at three different vial densities. Vial densities were 5, 10 and 15 flies per vial (10 replicate vials per variant, per sex).

### Lifespan assay

Vials were randomized and flies tipped into fresh food vials without anesthesia every 3-4 days. On these occasions, dead flies were counted and removed to prevent them from being tipped into the fresh food vials. Survivorship was scored at the time of tipping until all flies had died. Flies that escaped while tipping were censored. Thus, for each specific combination of genotype, sex, mating, and density the minimum number of flies was 50 and the total number of flies was 4800 before censoring. This factorial design enables us to quantify the effects of genotype, sex, social environment, and their interactions on phenotypic variation for lifespan.

### Statistical analyses

To compare the effects of sex, genotype, mating, and density on adult lifespan, we used a mixed model analysis of variance using restricted maximum likelihood (REML) estimates of the variance components (PROC GLIMMIX in SAS 9.2). Sex, DsGRP genotypes (DsGRP20♂ × DsGRP20♀, DsGRP57♂ × DsGRP57♀, DsGRP20♂ × DsGRP57♀, and DsGRP57♂ × DsGRP20♀,), mated status (non-mated/mated), density (5, 10 and 15 flies per vial) and their interactions were modelled as fixed factors and tested with F-statistics. For tests of fixed effects, we applied a Satterthwaite approximation to calculate the denominator degrees of freedom via the “ddfm=SAT” option in SAS. Vial was modelled as a random effect. Density was treated as a categorical factor as we did not necessarily expect linear relationships between density and longevity. Models were simplified by backward single term deletions (p ≤ 0.05). Significant interactions that included sex were explored by fitting the mixed model separately for each sex.

In our initial modelling, we used a four-level ‘genotype’ effect that includes the homozygous founder lines (DsGRP20 and DsGRP57) and both reciprocal F1 crosses between these lines. Subsequent contrasts between these four levels allowed us to test multiple genetic effects. First, we compared homozygous line differences to assess genetic differences in lifespan. Second, contrasts between the F1 and homozygous genotypes permitted a test for the effect of inbreeding. Third, contrasts between the two F1 crosses allowed us to test for a reciprocal cross effect that includes X chromosome genome influences. We present effect sizes as least square means and used Tukey’s HSD to correct for multiple testing.

## Results

After censoring 194 flies that escaped while being transferred to fresh holding vials (<5% of total flies), 4606 flies were available for analysis. Across the entire experiment, female-biased longevity was apparent. While female *D. serrata* lived on average 54 days (range 4 – 104 days), males lived an average of only 34 days (range 4-69 days). The final simplified linear model describing genetic and environmental influences on lifespan variation appears in Table 1. While the model provided statistical support for sex differences in lifespan in *D. serrata* (*Sex: F*_*1, 454*.*4*_ = 1798.3, *P* = 4.54^e-160^), males and females were influenced differently by genotype (*Sex × Genotype: F*_3,454.3_ = 64.6, *P =* 8.36^e-35^), which was also a significant main effect in the model (*Genotype: F*_*3,601*.*3*_ = 340.4, *P* = 3.83^e-129^). Here, three key results are of interest. First, reciprocal crossing did not affect the degree of sexual dimorphism with no lifespan differences found between the males of F1 genotypes (20♂x57♀) and (57♂ x 20♀) or between the females of these two F1 genotypes (Fig. 1). Second, males and females were affected by outcrossing in different ways. F1 females lived at least 17 days longer than homozygous parental line females and a similar degree of increase (∼ 40%) was observed in F1 males compared to parental line DsGRP20 males (Fig. 1). However, there was no difference in male lifespan between the F1s and parental line DsGRP57 (Fig. 1) consistent with a lack of any outcrossing effect. Third, genetic differences were also apparent between the two parental lines with both males and females from line DsGRP57 living between 14 and 7 days longer than males and females from line DsGRP20 respectively.

**Table 1.** *F*-tests of fixed effects for the reduced model examining the significance of contributions of sex, genotype, mating, and density to *D. serrata* lifespan.

| **Effect** | ***d.f.*** | ***F*** | ***P*** |
| --- | --- | --- | --- |
| *Sex* | 1, 454.4 | 1798.3 | 4.54e<sup>-160</sup> |
| *Genotype* | 3, 601.3 | 340.4 | 3.83e<sup>-129</sup> |
| *Sex × Genotype* | 3, 454.3 | 64.6 | 8.36e<sup>-35</sup> |
| *Density* | 2, 552.9 | 1.18 | 0.308 |
| *Genotype × Density* | 6, 552.7 | 0.84 | 0.539 |
| *Mating* | 1, 601.5 | 0.09 | 0.764 |
| *Genotype × Mating* | 3, 601.3 | 3.28 | 0.021 |
| *Density × Mating* | 2, 552.9 | 15.0 | 4.53e<sup>-07</sup> |
| *Genotype × Density × Mating* | 6, 552.8 | 2.45 | 0.024 |

**Figure 1.**
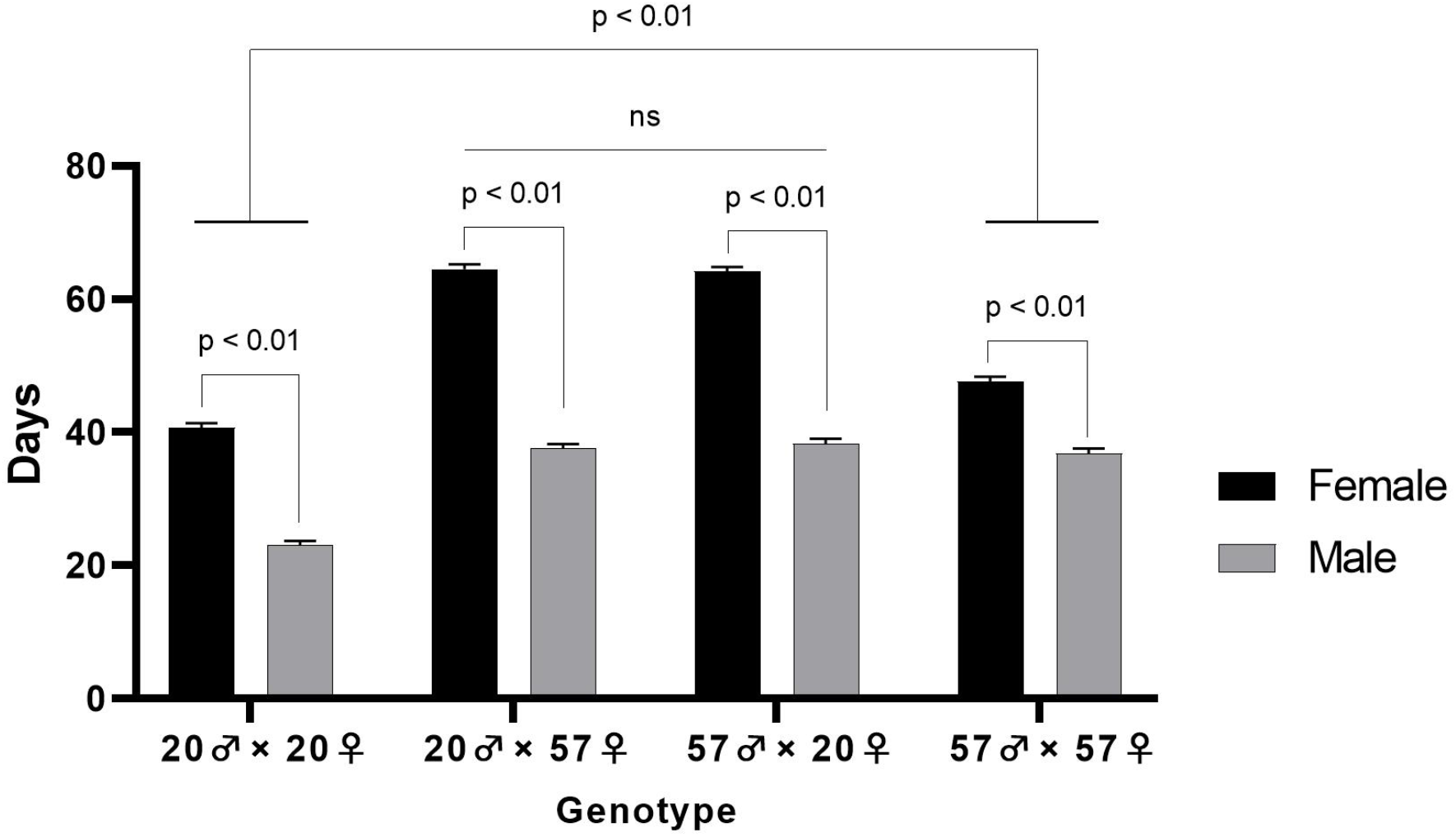
Genotype dependent effects on sex differences in lifespan in *D. serrata*. Sex differences in mean adult life span in the four genotypes resulting for our reciprocal cross between DsGRP20 and DsGRP57 (two parental lines plus alternate F1s). Bars represent the mean lifespan of each genotype pooled across the six density (low, medium, and high) × mating status (mated and non-mated) treatment combinations. Error bars represent 1 S. E.

Our analysis also indicated a genotype-by-environment interaction for lifespan. Genotype dependent effects were observed for both density and mating via a significant three-way interaction (Table 1: *Genotype × Density × Mating: F*_*6,552*.*8*_ = 2.45, *P* = 0.024). The social environmental effects underlying this significant interaction were, however, typically more subtle than the effects seen in the interaction between sex and genotype (Fig. 2) Considering this interaction further, post-hoc comparisons revealed significant differences between density and mating within only two of the four genotypes the parental (57♂x57♀) and the reciprocal F1 (20♂x57♀). For these genotypes, an effect of mating was detected but only in the low-density treatments, with the lifespan of mated flies on average, 6 days higher than unmated flies (Fig. 2).

**Figure 2.**
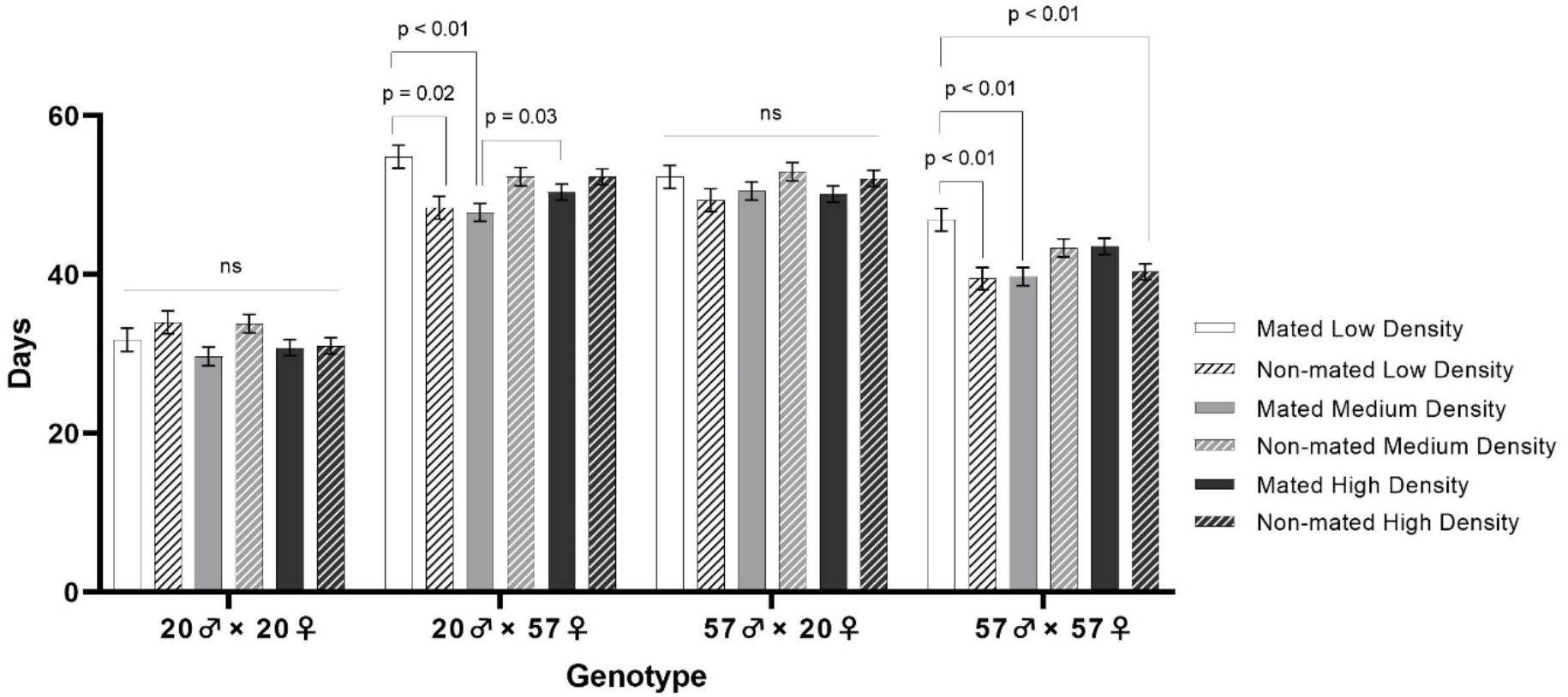
Genotype dependent effects of the social environment on lifespan in *D. serrata*. Shown are pooled adult, male and female lifespan for the homozygous founder lines *DsGRP20♂ × DsGRP20♀* and *DsGRP57♂ × DsGRP57♀*, as well as both reciprocal F1 crosses *DsGRP20♂ × DsGRP57♀* and *DsGRP57♂ × DsGRP20♀* between these lines. Each bar represents the mean of each genotype measured in one of six different density (low, medium, and high) × mating status (mated and non-mated) treatment combinations Error bars represent the 1 S.E. of the mean.

## Discussion

### Unguarded X and female-biased lifespan in D. serrata

All treatment combinations female *D. serrata* lived longer than males, a result consistent with a wide range of wild and captive species, where on average, the homogametic sex lives longer than its heterogametic counterpart (Xirocostas et al., 2020). Our result is also consistent with two previous studies of *Drosophila serrata* both of which indicate female-biased longevity (Robson et al., 2006, Wit et al., 2015). One prominent hypothesis for reduced male lifespan is the “unguarded X” hypothesis (Trivers, 1985). This hypothesis predicts that reduced male lifespan is a result of the unconditional expression of recessive deleterious alleles on the single X chromosome. To date, the few studies that have explicitly tested predictions arising from the unguarded X hypothesis, conducted in *Drosophila melanogaster* (Carazo et al., 2016, Sultanova et al., 2018, Brengdahl et al., 2018a) have produced inconsistent results.

Here, we used two inbred lines with differing lifespans to create outbred and reciprocal F1’s to test for reduced lifespan in males as predicted by the unguarded X hypothesis. Despite differences in inbred parental lifespan, we found no differences in lifespan between the outbred and reciprocal male F1’s that could be attributed to the accumulation of recessive deleterious mutations on the X chromosome as predicted by the unguarded X hypothesis (Fig. 1). Under the unguarded X hypothesis, outbred male F1 offspring of the shorter-lived maternal line inherit deleterious mutations on their X chromosome, resulting in lower lifespan than offspring from the longer-lived maternal line without recessive deleterious mutations on the X chromosome. Although the effects of recessive deleterious mutations may be underestimated in crosses between highly inbred lines due to higher expected levels of purging during the inbreeding process (Hedrick, 1994), similar to studies in *D. melanogaster* (Brengdahl et al., 2018a), the unguarded X hypothesis is appears to be insufficient to explain sexual dimorphism in *D. serrata* lifespan. Sex-specific differences in selection (Bonduriansky et al., 2008, Maklakov et al., 2009, Maklakov and Lummaa, 2013) could better explain the pattern of higher mortality in males and lifespan dimorphism observed in *D. serrata*. Alternative explanations that partly explain the patterns predicted by the unguarded X hypothesis and could be explored in future studies include sexually antagonistic genes and sex-specific expression patterns (Sultanova et al., 2018, Brengdahl et al., 2018a).

### Genotype-by-social environment interactions for lifespan

In addition to sex- and genotype-biased longevity, we also found interactions of genotype with mating and density, our two experimentally manipulated axes of social background. Across a range of taxa, sexual dimorphism is a result of complex relationships between environmental conditions and sex-specific reproductive costs (Lemaitre et al., 2020). Mean lifespans did not differ significantly between density treatments within genotypes (Fig. 2), even though large sex and genotype effects were detected. While we detected no *Genotype × Density* or *Sex × Density* interaction, there was a highly significant interaction between density and mating that appeared to be driven by a change in rank order lifespan between low and medium density, which was highest for low density in the mated treatment but lowest for the unmated treatment (Fig. 2). Survivorship experiments with high densities at the beginning can produce high mortality rates at young ages (Graves and Mueller, 1993), however we observed no such effect in our high density treatments.

In our study, mating had no effect on mean lifespan in *D. serrata*. While we did detect a significant *Genotype × Mating* interaction this can be explained by idiosyncratic effects of genotype on mating and density (Fig 1). Adverse effects of multiple mating on lifespan in *D. melanogaster* males have been reported in several studies, as have toxic effects of male accessory gland proteins on female fitness and lifespan (Fowler and Partridge, 1989, Chapman et al., 1995). In female *D. serrata*, continued male courtship and harassment also leads to decreased fitness in females (Chenoweth et al., 2015). Intermittent and short-term mating, as was the case in this study, could explain why mated and unmated flies have similar lifespans, except at low density in two genotypes where unmated flies lived on average 6 days longer. Though widespread, trade-offs between longevity and reproduction are hardly ubiquitous, can be highly plastic, and uncoupled under certain environmental or genetic conditions (Flatt, 2011).

## Conclusion

Here, we show that the pattern of sexual dimorphism in *D. serrata* is consistent with females living longer than males across all genotypes and treatments. As expected, outbred genotypes lived longer, and female lifespan was more adversely affected by inbreeding. However outbred male lifespan for the outbred F1 genotypes did not differ as expected from a cross between parental genotypes with significantly different lifespans. Overall, our findings converge with existing evidence to suggest that sex-specific selection largely drives the sexual dimorphism seen in lifespan (Bonduriansky et al., 2008, Maklakov et al., 2009, Maklakov and Lummaa, 2013), and that physiological differences resulting from strategies developed amongst sexes to maximize fitness can be independent of the effects of mating and/or density (Sultanova et al., 2020, Maklakov et al., 2017, Harvanek et al., 2017, Kimber and Chippindale, 2013, Ziehm et al., 2013, Vermeulen and Bijlsma, 2004a, Vermeulen and Bijlsma, 2004b). As the first study dissecting contributions of genetic background and social environment on lifespan in *D. serrata*, the robustness of these findings will no doubt be revealed by further testing effects on lifespan across different conditions. It is however reasonable to conclude that, based on a variety of studies across different taxa and Drosophila species, ageing in *D. serrata* is best viewed as a condition-dependent environmental modulation of a genetically determined trait.

## Acknowledgements

We thank Nicholas Appleton for assistance with fly work.

## Competing interests

The authors declare no competing financial interests.

## Author Contributions

All authors contributed to the planning of the experiments; V.P.N performed the experiments and the other authors assisted in analyses, interpretation, and writing the manuscript.

## Funding

This research was supported with funding provided by the University of Queensland. V.P.N. was supported by the QUEX Institute - a joint initiative of The University of Queensland and the University of Exeter.

## Notes

### Competing Interest Statement

The authors have declared no competing interest.

